# The role of neutrophil typing in homeostatic resilience protective model of inflammatory preconditioning

**DOI:** 10.1101/2023.08.17.553795

**Authors:** Liyuan Li, Xinfang Zhang, Hongmei Yu, Chao Yan, Jicheng Sun, Yan Qi, Yawei Gou, Mingming Zhang, Shengnan Wang, Haokun Li, Wenying Nie, Renhao Wang, Jiashan Jiang, Feng Gao, Xinyu Li, Qinghong Shi, Lina Song, Fang Wang, Xuesong Xu, Wei Sun

**Author notes:** Corresponding author:.(WS),.(XX), (FW). These authors contributed equally to the work (Liyuan Li, Xinfang Zhang, Hongmei Yu).

## Abstract

Sepsis is a systemic inflammatory response syndrome (SIRS) characterized by a dysregulated host response to infection that results in organ dysfunction. The response of neutrophils to early inflammation is critical, interferon regulatory factors 4 (IRF4) and interferon regulatory factors 5 (IRF5) are expressed in most cell types of the immune system, but the relationship between the role of IRF4 and IRF5 in preconditioning-protected septic peritonitis mouse model and neutrophils response remains unknown. In this study, we used an *E.coli*-induced septic peritonitis mouse model and zebrafish model to explore the relationship between neutrophil inflammation and IRF subtype changes in the process of septic peritonitis. Mechanistically, we found that the protein complex formed by polymorphonuclear neutrophils (PMN) and IRF5/MyD88 can secrete pro-inflammatory factors and directly kill invading bacteria.

**Author Summary:** Numerous animal and clinical sepsis studies have shown that markedly impaired recruitment of neutrophils to sites of infection and failure to clear bacteria contribute to excessive inflammation and increased mortality in sepsis. In our previous work, we established an *Escherichia coli* (*E.coli*) lethal septic peritonitis model and a preconditioning-protected septic peritonitis mouse model. In the early stage of septic peritonitis, a large number of PMN infiltrate the peritoneal cavity, which produces an inflammatory response to invasive bacteria. In this study, the switching process between pro-inflammatory and anti-inflammatory attracted our attention. Our research shows that IRF5-IRF4 regulatory axis leads to a PMN phenotype switch in sepsis and the expression of IRF5 and IRF4 was related to PMN pro-inflammatory (N1 type) and anti-inflammatory (N2 type) phenotypes, respectively.

## Introduction

Sepsis is defined as a life-threatening organ dysfunction caused by a dysregulated host response to infection[1]. During sepsis, inflammation and immunosuppression may occur sequentially or concurrently[2]. Bacterial infection in the abdominal cavity has always been one of the main causes of sepsis, which is known as intra-abdominal sepsis or septic peritonitis. Neutrophils, also known as polymorphonuclear (PMN) leukocytes, are important innate immune cells that can quickly migrate to the site of infection when inflammation occurs and become the host’s first line of defense against sepsis, acting as a “guardian”[3]. However, in the inflammatory response, PMN is a “double-edged sword”: with the development of the disease, overactivated PMNs release cytokines, triggering a cytokine storm, breaking the balance of pro-inflammatory and anti-inflammatory, thus further aggravating local tissue damage[4]. IRFs are a class of transcription factors with diverse biological activities in innate and adaptive immune responses and are determinants of the switch between pro-inflammatory and anti-inflammatory phenotypes of macrophages[5]. IRF4 and IRF5, as important members of the IRF transcription factor family, are expressed in most cell types of the immune system. Competitive binding of IRF5 and IRF4 to MyD88 is decisive for the polarization of macrophages into M1 or M2 phenotypes, which play a major role in pro-inflammatory and anti-inflammatory pathways, respectively[6–9].

In the early stage, when the research group established the “ *E. coli*-induced mouse septic peritonitis death model”, it was accidentally discovered that when we pre-injected a low dose of *E. coli* with 0.1×10^8^ colony forming units(CFUs)/mL into the mice intraperitoneally, after 2 h, and then injected a lethal dose of *E. coli* (3×10^8^ CFUs/mL), all the mice survived within 72 h (Fig. 1A) [10]. This suggested that small doses of *E. coli* stimulated the body to produce a pre-adaptation protection mechanism. In this study, by observing the expression of chemokine profiles at different time points in protective and lethal animal models, it was preliminarily determined that the correlation of IRF4/IRF5 with PMN pro-inflammatory and anti-inflammatory protein markers. We found that changes in the ratio of IRF4/IRF5 and its competitive binding with MyD88 are the key factors to distinguish pro-inflammatory and anti-inflammatory in PMNs.

**Figure 1.**
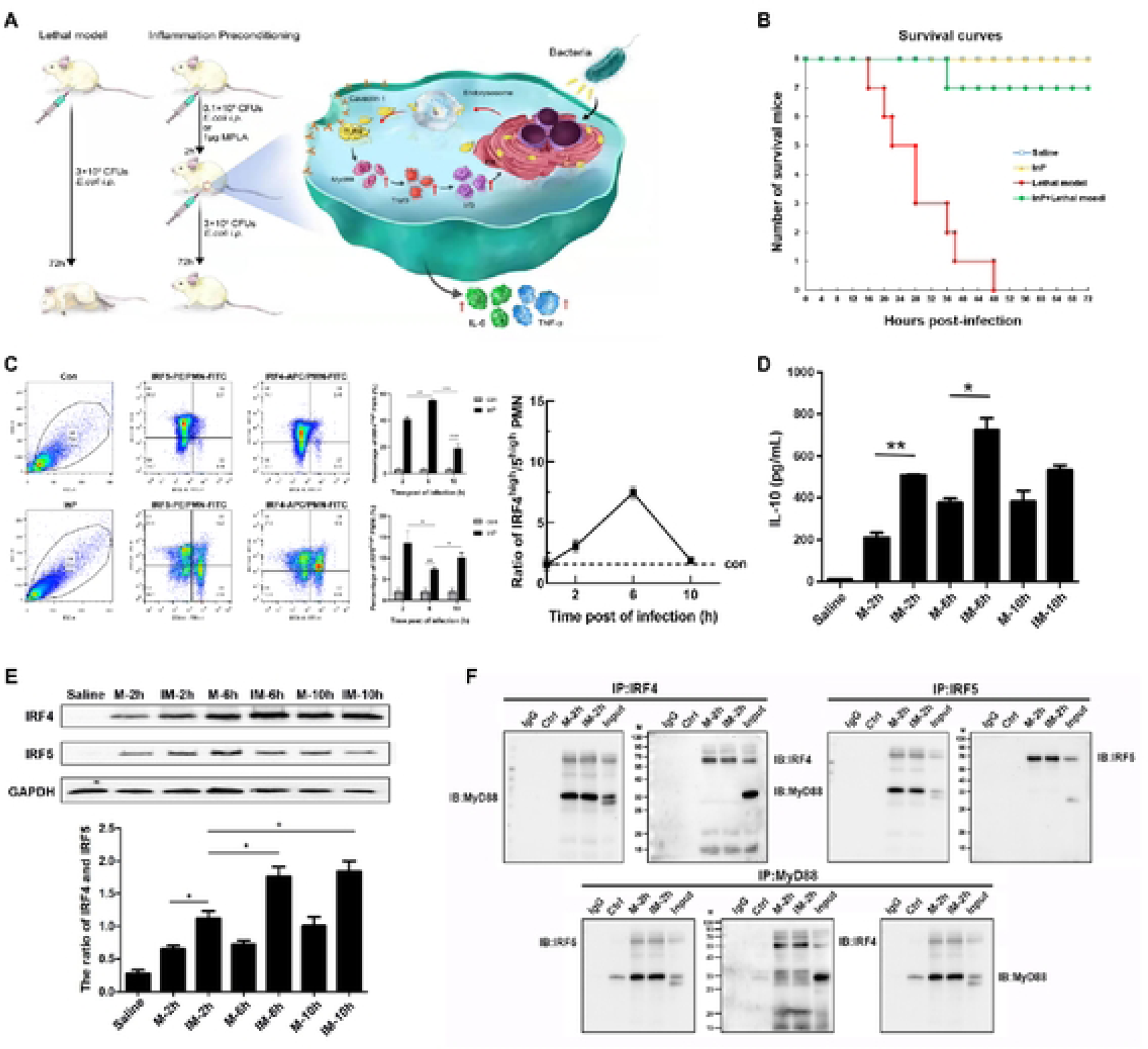
The increased ratio of IRF4/5 plays an important role in the mouse inflammatory preconditioning model. (A) Schematic diagram of mouse *E. coli* inflammation model[10]. InP group was injected intraperitoncally with 0.1×10^8^ CFUs/mL *E. coli,* Lethal model group was only injected with 3×10^8^CFU/mL *E. coli,* IM group was first injected with 0.1×10^8^CFUs/mL of *E. coli,* and 2 h later injected with 3×10^8^ CFUs/m L of *E. coli (n=* 10/group). (B) Survival curve of mice within 72 h. (C) The expression levels of IRF4 and IRF5 in peritoneal PMNs after inflammatory preconditioning were detected by flow cytomctry. (D) Statistical chart of IL-10 Model group and IM group at 2, 6 and 10 h after i.p. of *E coli.* (F) The interaction between IRF4 and IRF5 with MyD88 by Co-IP and reciprocal Co-IP experiments.

## Results

### Anti-inflammatory protection of PMNs mediated by elevated IRF4/IRF5 ratio in mouse sepsis model and inflammatory preconditioning model

To study the protective effect of preconditioning mice before injecting a lethal dose of *E. coli*, we established a model of bacterial peritonitis induced by intraperitoneal injection (i.p.) of *E. coli* in mice. The experiment was divided into four groups: blank control group (Saline), *E. coli* lethal model group (Lethal model, Model), independent inflammatory preconditioning group (InP) and inflammatory preconditioning protection group (InP+Lethal model, IM). The survival of the mice showed that compared with the control group, the mice in the model group died from 16 h, and the mortality rate was 100% within 48 h; in the IM group, only one mouse died within 36 h, and the mouse survival rate reached 87.5% within 72 h. No mouse died in InP group within 72 h (Fig. 1B). The results showed that pretreatment with low doses of *E. coli* protected the mice from death resulting from subsequent lethal doses of *E. coli* infection.

Studies have shown that IRF5 and IRF4 in microglia are associated with pro-inflammatory and anti-inflammatory responses, respectively, after stroke[8]. In order to verify whether this protective effect of preconditioning is also related to IRF5 and IRF4 during sepsis infection in the mice, we detected the expression levels of IRF4 and IRF5 in peritoneal PMNs by flow cytometry. The results showed that the expression levels of IRF4 and IRF5 increased in PMNs of the InP group compared with the control group (Fig. 1C). the highest expression level of IRF4 was observed at 6 h, while the lowest expression level of IRF5 was observed at this time. Further analysis of changes in peripheral blood cytokine expression levels, the results showed that the levels of the cytokine IL-10 in the IM group increased significantly with the duration of infection and reached the peak at 6 h. In different stages of inflammation (2, 6, 10 h), the expression level of IL-10 in the peripheral blood of IM group was higher than that of the Model group without pretreatment (Fig. 1D). The changes of IRF4 and IRF5 protein expression in the peritoneal lavage fluid cells (PLCs) of mice in Model group (M) and IM group were detected by western blot at corresponding time points. Compared with the control group (Saline), the expression of IRF4 and IRF5 proteins in the M group increased; In the IM group, the expression of IRF4 protein gradually increased, the expression of IRF5 protein gradually decreased and the ratio of IRF4/IRF5 showed a significant upward trend (Fig. 1E, *P*<0.05). The above results indicated that pretreatment before i.p. of a lethal dose of *E.coli* in mice may increase the ratio of IRF4/5, promote the secretion and expression of IL-10, and play an anti-inflammatory protective effect.

The competitive binding of IRF4 and IRF5 to MyD88 is a decisive factor in the pro-inflammatory or anti-inflammatory phenotype of macrophages. In order to verify the role of IRF4 and IRF5 in the mouse inflammatory preconditioning model, we detected the co-expression of IRF4, IRF5 and MyD88 in the PLCs of the Model group and the IM group in the early stage of inflammation (2 h after i.p injection the lethal dose of *E.coli* in mice) by Co-Immunoprecipitation experiment (Co-IP). The results show that MyD88 is significantly enriched when IP-IRF4 is applied; at the same time, MyD88 is also significantly expressed when IP-IRF5 is applied; the enrichment of IRF4 and IRF5 proteins also appeared in Model and IM groups when IP-MyD88 (Fig. 1F). The above results indicated that IRF4 and IRF5 were co-expressed with MyD88 in mouse sepsis model and inflammatory preconditioning model.

### Both IRF4^+^IRF5^−^ and IRF4^+^IRF5^+^PMNs are involved in the inflammatory process, but IRF5^+^ PMNs play a key pro-inflammatory role in the differentiation of inflammatory phenotypes

To further determine the role of IRF4 and IRF5 in the mouse model of inflammatory preconditioning, we took PLCs for flow cytometry detection, and analyzed the expression of IRF4 and IRF5 in PMNs under different infection states (Fig. 2A). PMN cell populations were determined by double positivity for Ly6G^+^ and CD11b^+^ by flow cytometry. At 6 h after modeling, 20.4% of PMNs in the IM group expressed IRF5 and 99.1% of PMNs expressed IRF4. These results indicated that PMN expressed high levels of IRF4 and low levels of IRF5 after inflammatory preconditioning (Fig. 2B). Further analysis found that the ratio of IRF5^+^ PMN was significantly increased in different infection states, especially in the Model group. In the IM group, although the same amount of lethal *E. coli* was also injected, the ratio of IRF5^+^ PMN was significantly lower than that of the Model group at 2 h to 6 h after injection. IRF4^+^ PMN had no significant change except in the InP group alone. the ratio of IRF5^+^ and IRF4^+^ PMN was highest at 2 h after *E.coli* infection, and the change trend of IRF5^+^/IRF4^+^ PMN was consistent with the change of IRF5^+^ PMN (Fig. 2C). It shows that IRF5^+^ PMN is positively correlated with the degree of infection, while the change of IRF4^+^ PMN cannot fully reflect the degree of infection.

**Figure 2.**
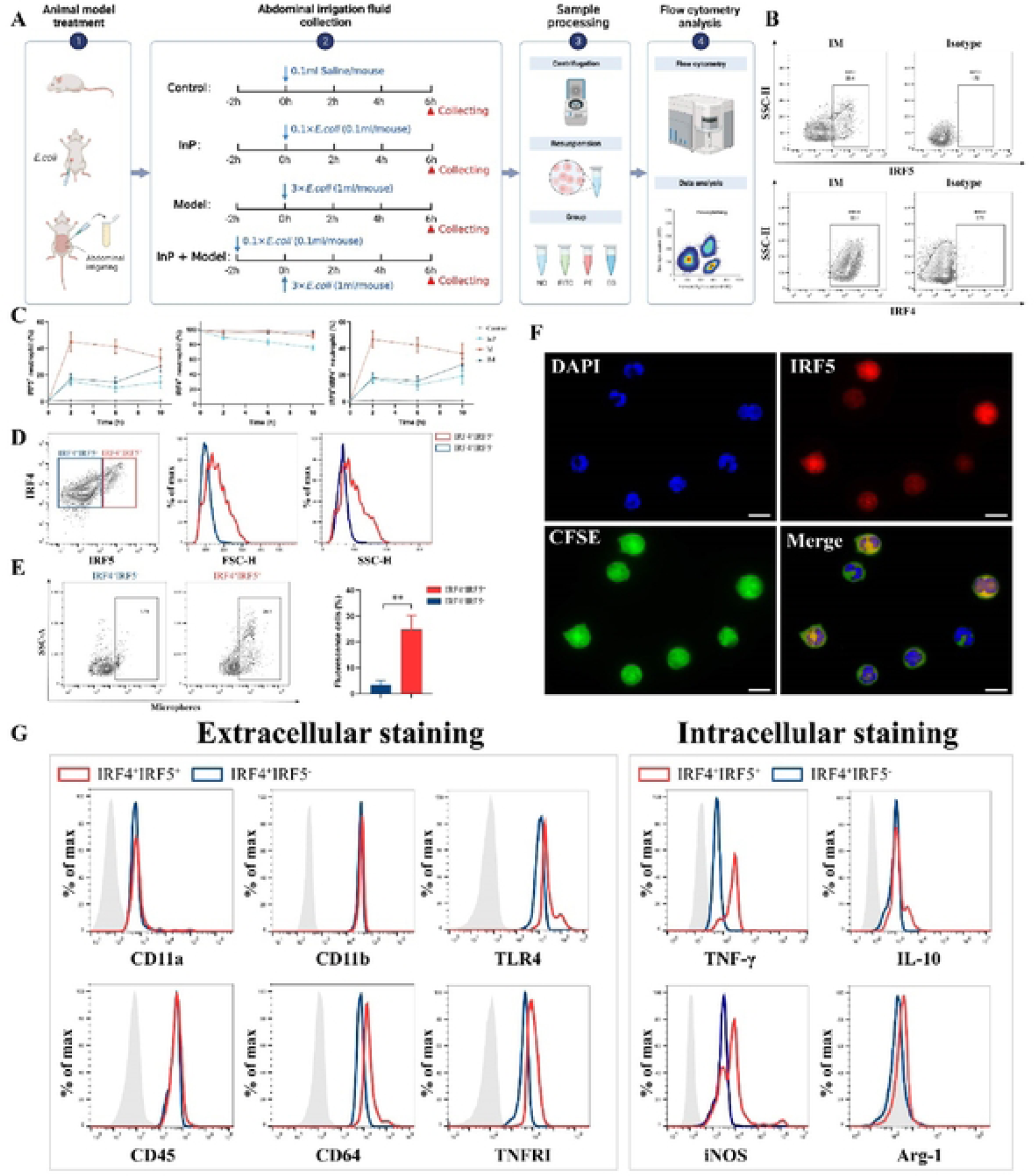
The expression of IRF4 and IRF5 and the changes of related inflammatory factors in different infection states. (A) Schematic diagram of flow cytometry experiment. (B) The expression levels of IRF4 and IRF5 in neutrophils in IM group, and Isotype is the peer control group. (C) The amount of IRF5^+^PMN and IRF4^+^PMN in mouse peritoneal lavage solution. (D) based on the differential expression oflRF4 and IRF5 in PLCs of mice 6 h after infection. (E) Proportion of 1Rf4^+^IRF5^−^PMN and 1Rf4^+^1Rf5^+^PMN. (F) IRF5 expression in neutrophils was detected using IF. DAPI-stained nuclei are blue, CFSE-stained intracellular proteins are yellow-green, and IRF5 protein is stained red, the scale bar=20 µm. (G) Changes of inflammation-related factors in IRF4^+^IRF5·PMN and IRF4^+^IRF5^+^PMN cell populations.

The transformation of PMN pro-inflammatory and anti-inflammatory phenotypes is particularly important to reflect the degree of inflammatory infection, so we identified PMN subsets by Ly6G and CD11b expression, then according to the difference in the expression of IRF4 and IRF5, PMNs were identified as two distinct cell populations, namely IRF4^+^ IRF5^−^ and IRF4^+^ IRF5^+^ (Fig. 2D), the proportions of IRF5^−^ and IRF5^+^ were 1.79% and 24.8%, respectively, and the difference between the two groups of cells was statistically significant (Fig. 2E). Compared with IRF4^+^ IRF5^−^ PMN, the IRF4^+^ IRF5^+^ PMN population had larger cell volume and more intracellular granules (Fig.2E). Immunofluorescence staining (IF) results showed that the fluorescence intensity of the IRF5^+^ PMN group was significantly enhanced, the cell volume was larger, the intracellular protein content was more, the nucleus lobulation was more obvious, and the intracellular granule density increased significantly (Fig. 2F). It shows that IRF4^+^ IRF5^+^ PMNs are more mature than IRF4^+^ IRF5^−^ PMNs. Flow cytometry results showed that both IRF4^+^ IRF5^−^ PMN and IRF4^+^ IRF5^+^ PMN highly expressed inflammatory factors such as CD11a, CD11b, CD45, Arg-1 and IL-10. However, IRF4^+^ IRF5^+^ PMNs expressed higher levels of CD64, TLR4, and TNFR1, produced more IFN-γ and iNOS, and also had stronger phagocytic activity and a typical pro-inflammatory phenotype (Fig. 2G, Supplementary Fig. S1). The above results indicate that the IRF5^+^ PMN subset mediates the activation of pro-inflammatory cells and promotes the occurrence of inflammation.

### Inflammatory preconditioning maintains immune system function by activating PMN aggregation, chemotaxis, and number in both LPS-induced inflammation model and *E. coli* infection visualized zebrafish model

Zebrafish is a model animal which can quickly and efficiently construct tissue cell-specific gene editing, and zebrafish has the characteristics of real-time and visual observation of PMN migration and aggregation, and can detect quantitative changes of fluorescent bacteria in vivo[11]. We established an inflammatory lethal model by using wild-type (WT) zebrafish (3 dpf) soaked in lipopolysaccharide (LPS) and observed and statistically analyzed the survival of zebrafish (Fig. 3A&B). The survival results show that with the increase of LPS concentration, the mortality rate of zebrafish larvae increased and the survival time shortened (Fig. 3B). Compared with the control group, with the increase of LPS concentration (10-200 μg/mL), the inflammatory symptoms of zebrafish larvae gradually aggravated (Fig. 3C). However, 10 μg/mL LPS treatment only caused a slight inflammatory reaction in larva, and did not cause death of juvenile fish. Therefore, we chose 10 μg/mL LPS as the preconditioning dose for subsequent experiments and established the zebrafish InP model, chosing100 μg/mL LPS as the lethal concentration. The results showed that compared with the model group, the survival rate of larva in the IM group was about 90%, the survival time was significantly prolonged, indicating that low-concentration LPS pre-soaking can make zebrafish larvae acquire the ability to resist the subsequent lethal dose of LPS infection, and the experimental results are consistent with the experimental results of mice (Fig. 3D). Compared with the model group, it was found that in the IM group, the spine of the larvae did not bend, the light transmittance increased, and the damage and retraction of the tail also improved through morphological observation (Fig. 3E). qPCR results were consistent with the mouse sepsis and inflammatory preconditioning model, that is, IRF4 and IRF5 mediated the inflammatory response of PMN (Supplementary Fig. S2).

**Figure 3.**
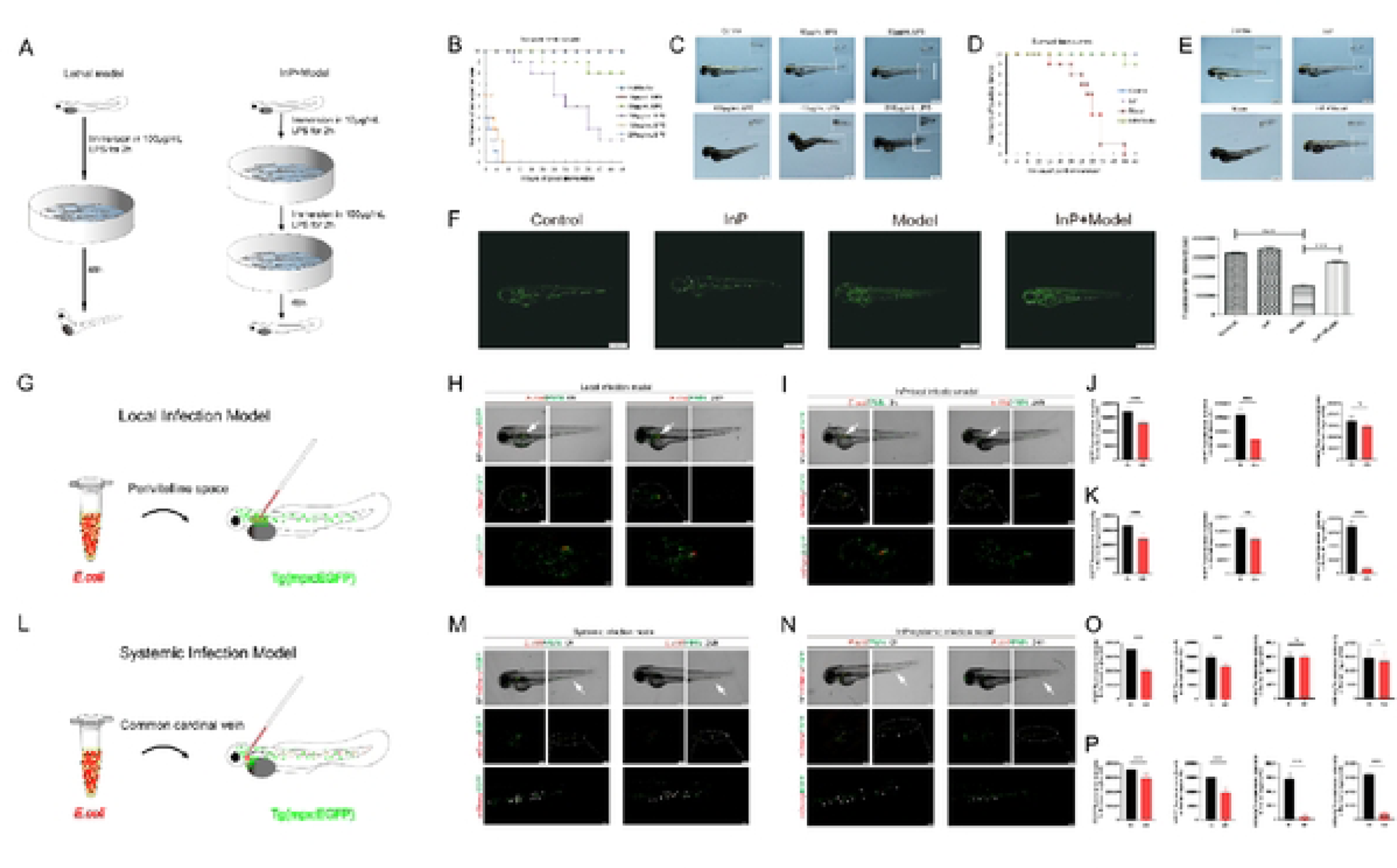
Establishment of LPS-induced zebrafish inflammation model and fluorescent *E. coli* zebrafish inflammation model. (A) Schematic diagram of LPS pretreatment 7.cbrafish model. the InP group was soaked in 10µg/mL LPS, the Model group was soaked in 100µg/mL LPS, and the InP^+^Model group was first soaked in 10µg/mL LPS for 2 h, and then soaked in 100µg/mL LPS soaked *(n=*I0/group). (B, C) Survival curves and body surface changes of larvae soaked in different concentrations of LPS. The zebrafish larvae were soaked in LPS for 2 h and then transferred to fresh Holt buffer solution for culture to observe their survival. (D, E) Survival curves and surface changes of larvae in LPS-induced inflammatory lethal model and neutrophils in zebrafish *(n=*I0/group). (G-K) Establishment of a visual model of local infection in zebrafish. (L-P) Establishment of a visual model of systemic infection in zebrafish. 3 x 10^2^ CFUs *mCherry-E.coli* were injected in local or common cardinal vein of Tg(mpx: EGFP) zebrafish larvae. Zebrafish larvae of InP+local infection model group/InP+systemic infection model group were first soaked in 1 mL 0.1 x 10^8^ CFUs/mL mCherry-E.co/i solution, then injected about 3×102 CFUs *mCherry-E.coli.* Arrows indicate primary or metastatic *E.coli* sites. Dashed lines circle and amplify the indicated regions. (scale bars = 200 µm; in amplified fields= 100 µm).

During infection and subsequent release of pro-inflammatory mediators, neutrophils migrate toward the infection site, then recognize and engulf the pathogen[12]. We used Tg(mpx:EGFP) transgenic zebrafish to observe the changes of neutrophils after LPS inflammatory pretreatment. The experimental results showed that, compared with the control group, the number of neutrophils in the model group was significantly reduced, while the number of neutrophils in the IM group was significantly higher than that in the model group (Fig. 3F). Next, we established bacterial local infection and systemic infection visualization models by injecting fluorescent *E. coli* into the yolk sac space and the common cardinal vein, respectively (Fig. 3G&L). In the systemic infection model established by injecting mCherry-*E. coli* into the common cardinal vein, *E. coli* will diffuse throughout the larvae with the blood circulation, and *E. coli* labeled with red fluorescence can be clearly observed in the whole body, tail, heart and yolk sac, etc. The experimental results showed that, no matter in the *E. coli* local infection model or the systemic infection model, PMNs in juvenile fish were chemotaxis and aggregated within 0-30 min after injection of large doses of mCherry-*E.coli*. In the localized infection model, PMNs chemotaxis and accumulate into the yolk sac space (Fig. 3H&I), and in the systemic infection model, PMNs were chemotaxis and accumulated in common veins (Fig. 3M&N). At 24 h after injection, in the local infection model, *E.coli* in the yolk sac space still existed in large quantities, and there was no significant difference in *E.coli* content compared with the group injected at 0 h (*P* >0.05, vs local infection model/0 h), while the number of PMNs was significantly reduced (*P*< 0.001, vs local infection model/0 h, Fig. 3H&J); similar results were shown in the systemic infection model (Fig. 3M&O).

In the local infection model, the *E.coli* content in the yolk sac space of juvenile fish significantly decreased 24 h after injection of lethal dose of *E.coli* in the InP+local infection model group (*P*<0.001, vs local infection model/24 h, Fig. 3I&K), and the number of PMNs in juvenile fish was significantly increased (*P*<0.001, vs local infection model/24 h, Fig. 3I&K). Similar results were shown in the systemic infection model: InP+systemic infection model group had a significantly lower *E.coli* content (*P*<0.001, vs systemic infection model/24 h, Fig. 3N&P) and a significantly higher PMN number in juvenile fish (*P*<0.001, vs systemic infection model/24 h, Fig. 3N&P).

### Potential application of zebrafish bacterial infection visualization model and zebrafish inflammation pre-conditioning model in the treatment of clinical bacterial infection

We analyzed the data from the China Antimicrobial Resistance Surveillance System in 2021. The top three bacteria in the list were *E.coli*, *Klebsiella pneumoniae*, and *Staphylococcus aureus*, which served as the basis for selecting these three bacteria to establish the model (Fig. 4A and Supplementary Table1). Further, we collected and analyzed the bacterial drug sensitivity test results of inpatients and outpatients in the China-Japanese Union Hospital of Jilin University in 2021. The data showed that the resistance rate of methicillin resistant *Staphylococcus aureus* (MRSA) to gentamicin, rifampicin, levofloxacin, and clindamycin was higher than that of *Staphylococcus aureus* (Fig. 4B and Supplementary Table2). The susceptibility of *E.coli* to ampicillin, cefazolin, cefuroxime, ceftriaxone, ciprofloxacin and levofloxacin was less than 50%, while the susceptibility to carbapenems was higher. The resistance rate of *Klebsiella pneumoniae* to cefazolin, cefuroxime and ceftriaxone was about 20% higher than other antibiotics (Fig. 4C and Supplementary Table3). Therefore, we selected six kinds of drug-resistant and non-drug-resistant *E.coli*, *Klebsiella pneumoniae*, and *Staphylococcus aureus* isolated from blood samples of patients with clinical bacteremia as following up test strains.

**Figure 4.**
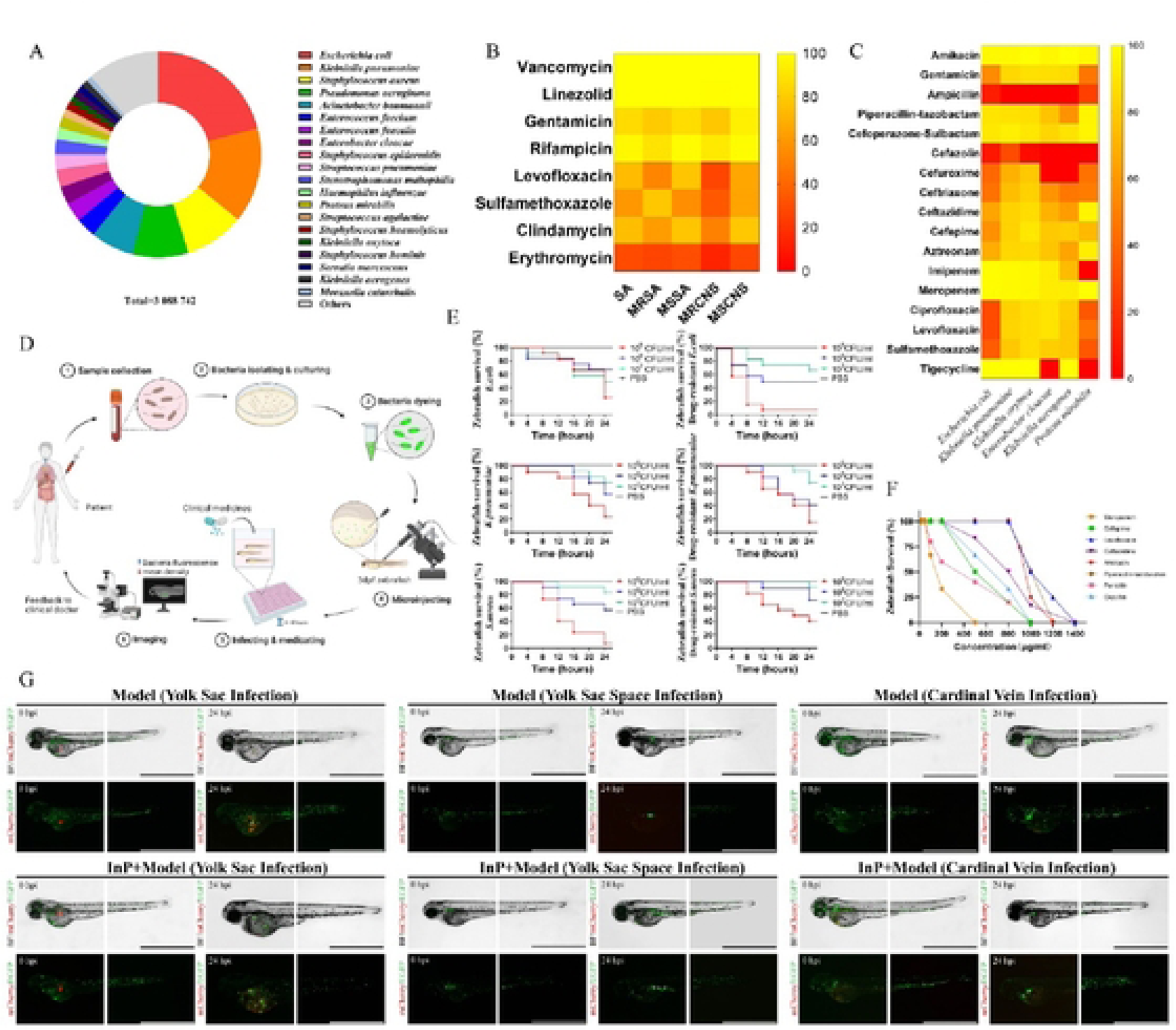
Establishment of zebrafish embryo infection model and drug screening platform. (A) Distribution characteristics of 3,088,742 strains of bacteria in China. (B) Susceptibility of Staphylococcus species to antimicrobial agents. SA, *Staphylococcus aureus;* **MRSA,** *Methicillin-Methicillin-resistant coagulase-negative staphylococci;* MSCNS, *Methicillin-sensitive coagulase-negative staphylococci.* (C) Susceptibility of Enterobacteriaceae species to antimicrobial agents. (D) Schematic representation of the in vivo drug-screening setup in the zebrafish bacterial infection model. (E) The survival curve of zebrafish infected with different clinical pathogens, the bacterial concentration of the subsequent infection model was 10^9^ CFUs/mL. (F) Drug toxicity test to determine the maximum drug concentration that can be tolerated by zebrafish. (G) Establishment of inflammatory-preconditioning models for local and systemic infections in zebrafish using clinical pathogenic strains. scale bar= 1000 *µm*.

To simulate the progress of clinical bacterial infection and explore the potential application of zebrafish bacterial infection model in clinical treatment, the zebrafish antibiotic screening platform was established. Firstly, collecting blood samples from patients. Secondly, isolating and culturing bacteria. Thirdly, dyeing the bacteria. Then, using the microinjection equipment to inject bacteria into zebrafish (3 dpf), and then immersing in antibiotics in 96-well plates. Observing the mean fluorescence intensity changes of bacteria in zebrafish for 0-4 h, so as to select effective antibiotics and feed back to doctors (Fig. 4D). In order to establish the antibiotic screening platform in vivo, we drew the survival curves of zebrafish infected with six kinds of bacteria and determined the bacterial concentration for subsequent infection models (Fig. 4E). In addition, a drug toxicity test was conducted to determine the maximum tolerable antibiotic concentration of zebrafish larvae. The result indicated that the safe concentration of antibiotics was 50 μg/ml for zebrafish (Fig. 4F). After transformation with clinically isolated and cultured strains, the systemic infection model and local infection model of zebrafish were established by injecting yolk sac, yolk sac space, and cardinal vein, respectively. The results showed that bacteria did not diffuse in zebrafish within 24 hours after injection of the yolk sac space, but after injection of the cardinal vein and yolk sac, bacterial diffusion, especially in the tail vein, could be seen. In addition, the bacterial content in the InP+Model group was lower than that in the Model group 24 hours after infection over time, indicating that the bactericidal ability of the InP+Model group was significantly stronger than that of the Model group (Figure 4G).

To establish a drug screening platform in vivo, 3dpf zebrafish larvae were used to observe the changes of mean fluorescence intensity after bacterial dyeing. It can be seen intuitively that, compared with the control group, the mean fluorescence intensity of the three kinds of bacteria has decreased to a certain extent with time (Fig. 5A), and the survival rates have improved (Fig. 5C). These results indicated that the antibiotics have certain effects on bacterial therapy. However, for drug-resistant bacteria, the results showed that levofloxacin, ceftazidime, ampicillin and oxacillin had poor efficacy in the treatment of bacteria. Compared with the control group, the mean fluorescence intensity was unchanged basically, and the survival rate was reduced to a certain extent (Fig. 5B and 5C). These results indicated that a visualized antibiotic screening platform for bacterial infection in vivo has been successfully established.

**Figure 5.**
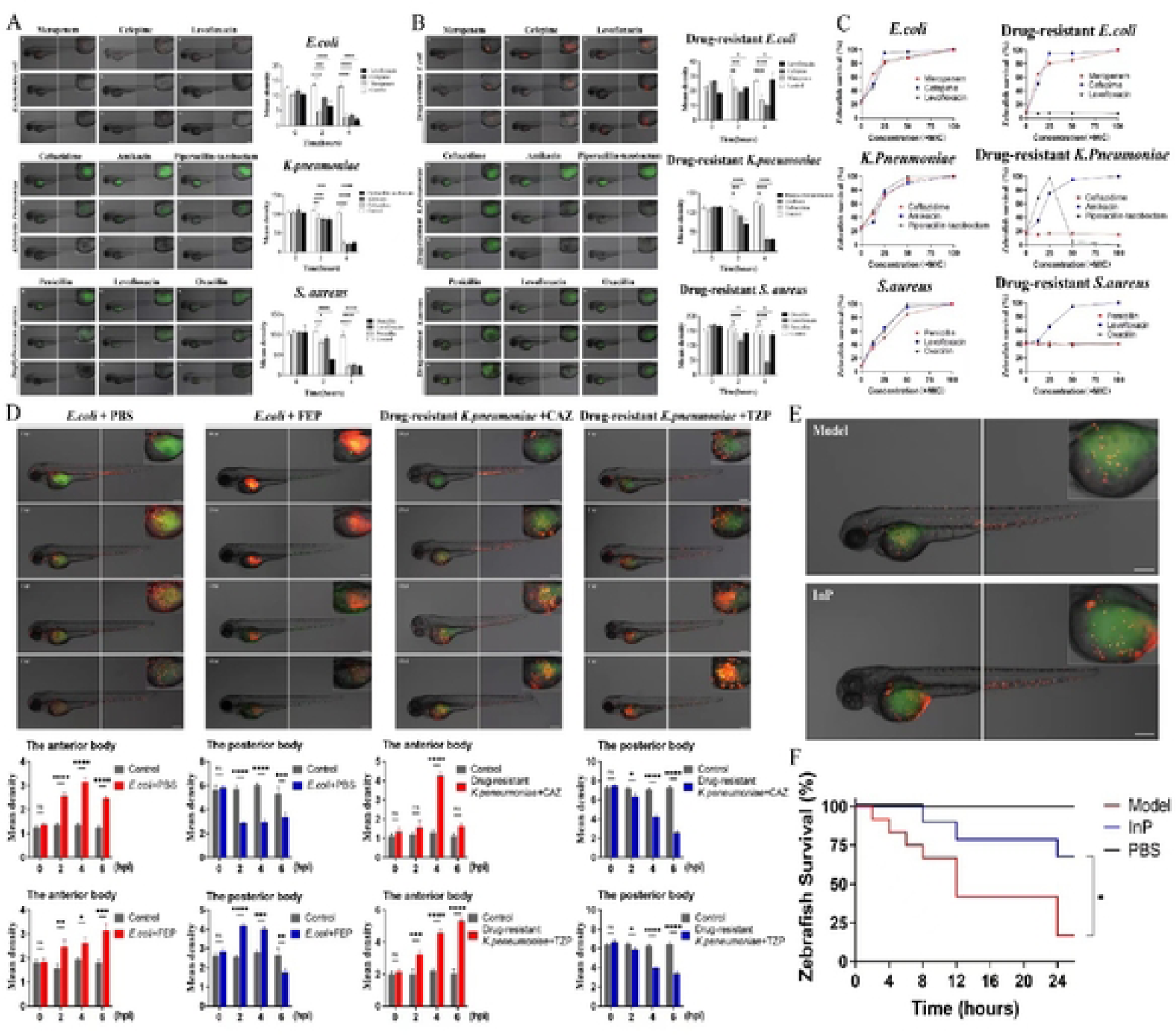
Zebrafish bacterial infection visualization model and inflammation pre-conditioning model can be used for the treatment of clinical bacterial infection. (A, 8) Clinical isolates of drug-resistant strains *E.coli, Klebsiella pneumoniae,* as well as *Staphylococcus aureus* and their corresponding drug-resistant strains were subjected to in vivo drug screening in zebrafish along with graphs of mean fluorescence intensity. Mean density= RawlntDent /Area. (C) Zebrafish survival changes before and after drug treatment. Survival was scored 4 dpi. (D) Aggregation of neutrophils in 6 h after infection of zebrafish with fluorescent markers of neutrophils and quantification of the mean fluorescence intensity of neutrophils at different sites. (E) Effects of inflammation pre-conditioning on neutrophils in lethally infected zebrafish. (F) The change of zebrafish survival rate by inflammatory pre-conditioning. Statistical significance was determined by Log-rank (Mantel-Cox) lest (*P<0.05). Each treatment group consisted of 12-30 zebrafish. Scale bars = 200 µm.

As a note, to observe the interaction between neutrophils and the host in the process of infection, Tg (LysC: DsRed2) and Tg (mpx: EGFP) with red and green fluorescent labeled neutrophils zebrafish were used, respectively. The results suggested that neutrophils gradually migrated from the hematopoietic tissue of the tail to the infected location within 2-4 h after infection, and the mean fluorescence intensity of neutrophils at the infected location was gradually increased, indicating that neutrophils play a critical role in bacteria elimination (Fig. 5D). Additionally, we established a zebrafish inflammatory pre-conditioning model. Compared with the model group, the finding indicated that the inflammatory pre-conditioning model (Low+high dose) group had more neutrophil aggregation (Fig. 5E), and the survival rate was significantly higher than the model (high dose) group (Fig. 5F). Therefore, we concluded that the zebrafish bacterial infection visualization model and the inflammatory pre-conditioning model are of significant in the treatment of clinical bacterial infections.

## Discussion

The pathophysiological mechanism of sepsis is complex and is a highly heterogeneous syndrome in which excessive inflammation and immunosuppression are intertwined in the development of sepsis[13]. The key proinflammatory responses in sepsis include activation of the complement system, coagulation system, vascular endothelium, neutrophils, and platelets[14]. sepsis is an inflammatory response disease, changes in pro- and anti-inflammatory phenotypes determine disease progression and correlate with patient outcomes[15], when sepsis patients require intensive care unit admission, one third of patients do not survive beyond 30 days[16, 17], moreover, the degree and duration of the inflammatory response depends on the degree and persistence of the threat (e.g. pathogen load) [18]. Animal models suggest that depletion of pro-inflammatory cytokines such as TNF, IL-1β, IL-12, and IL-18 confers robust protection against organ damage and death[19].

Neutrophils, as the most abundant cell type among circulating leukocytes, play a key role in sepsis-induced inflammatory pathophysiology and immune dysregulation[20–24]. However, neutrophils have a short half-life of only 18-19 hours. Neutrophils exhibit several hallmarks of immunocompromise in sepsis, Gram-negative bacteria, such as *E. coli*, which express cell-surface LPS, through a TLR4-induced signaling cascade activates innate lymphocytes to produce IL-17A, which triggers an increase in plasma granulocyte colony-stimulating factor [25]. Studies have shown that under septic insult, the spontaneous apoptosis of neutrophils is inhibited and the lifespan of neutrophils is increased[26]. Kinase activity is impaired in neutrophils from septic patients compared with uninfected critically ill patients, further suggesting an immunosuppressive phenotype of neutrophils [27]. Single-cell RNA-sequencing analysis reveals neutrophil heterogeneity, in the inflammatory state, neutrophils are divided into different populations, and the number ratio, functional characteristics, transcription factor expression and population transformation pathway of each population have obvious changes [28]. The latest study shows that two neutrophil-specific subsets, CD10^−^CD64^+^PD-L1^+^ neutrophils and CD10^−^CD64^+^CD16^low/−^CD123^+^ immature neutrophils, were identified by mass spectrometry and conventional flow cytometry, and may be a key factor driving the immune signature of sepsis[29].

IRF is a class of transcription factors with multiple biological activities in innate and adaptive immune responses, and is a decisive factor for the pro-inflammatory and anti-inflammatory phenotypes of macrophages[30]. In the study of macrophage polarization, the competitive binding of IRF4 to IRF5 MyD88 is a decisive factor in macrophage polarization towards the M1 or M2 phenotype, it was found that M1 type macrophages have the ability to promote inflammation and antigen presentation, The highly expressed IRF5 in M1 macrophages can bind to MyD88, induce the release of a large number of pro-inflammatory cytokines, such as TNF-α, IL-6, and IL-12, etc., and at the same time inhibit the production of anti-inflammatory cytokines [31, 32]. In M2 macrophages, the expression of IRF5 is low, leading to the production of a large number of anti-inflammatory cytokines such as IL-4, and IL-1[33–35]. In microglia, the expression of IRF5 and IRF4 are associated with pro-inflammatory and anti-inflammatory responses, respectively[8]. Down-regulation of IRF5 signaling by interfering RNA (siRNA) or conditional knockout (CKO) techniques resulted in increased IRF4 expression, enhanced M2 activation, quenched proinflammatory responses, and improved stroke outcomes, whereas down-regulation of IRF4 led to increased IRF5 expression, enhanced M1 activation, exacerbated proinflammatory responses, and worse functional recovery. Up-regulation of IRF4 or IRF5 by lentivirus induced similar results[8]. In hyperinflammatory diseases such as arthritis and systemic lupus erythematosus, IRF5 in PMN releases pro-inflammatory factors such as IL-1 through activation of TLR3/7, so that PMN can play a pro-inflammatory role[36]; IRF4 protein can significantly inhibit the early process of PMN-mediated severe asthma model, and affect the differentiation of Th17 cells in the subsequent disease process, so that PMN can play an anti-inflammatory role[37]. The above studies suggest that IRF is involved in the pro-inflammatory and anti-inflammatory process of PMN, and is associated with TLR subtypes. At the same time, MyD88 exists in the endoplasmic reticulum, endosome and cell membrane of PMN, which binds to the cell membrane and intracellular TLR subtypes respectively, thereby activating different signaling pathways[38, 39].

Inflammatory preconditioning can avoid the occurrence of severe sepsis, reducing the rate of severe sepsis, and small doses of pretreatment are expected to improve patient survival[40, 41]. We deeply studied septic peritonitis and found that InP plays a pro-inflammatory role in the early stage of inflammation and an anti-inflammatory effect in the later stage of inflammation by activating the IRF pathway. Septic peritonitis essentially initiates the complex process of host pro-inflammatory and anti-inflammatory. Our previous results showed that in the MPLA-induced inflammatory preconditioning model, the expression of IRF5 in mTLR9^+^ PMNs ranged from high to low over time. It is preliminarily shown that in the inflammatory preconditioning model, neutrophils first play a pro-inflammatory role, and then the pro-inflammatory role gradually weakens to avoid excessive inflammation in the body and thus play a protective role. It is speculated that after MPLA induces inflammatory preconditioning, IRF5/MyD88 in PMN forms a protein complex, secretes pro-inflammatory factors such as TNF-α, directly kills the bacteria that subsequently invade the body, and plays the role of body guards. PMNs at this time were named “type 1 pro-inflammatory neutrophils (N1 type)”. Then, with the occurrence of inflammatory response process, IRF4/MyD88 in PMN forms a protein complex, and releases anti-inflammatory factors IL-10 and TGF-β to play a role in inhibiting inflammatory response, thereby inhibiting the occurrence and development of subsequent septic peritonitis and avoiding the occurrence of excessive inflammatory response, we named PMN at this time as “type 2 anti-inflammatory neutrophils (N2 type)”. Our research group used the inflammatory preconditioning protection model, the mouse model of septic peritonitis and the zebrafish model to explored the molecular mechanism of pro-inflammatory and anti-inflammatory pathways. Our study found expression of IRF5 and IRF4 was related to pro- and anti-inflammatory responses, respectively. These experiments suggested that IRF5 and IRF4 signaling inhibit each other and that the manipulation of the IRF5-IRF4 regulatory axis leads to a PMN phenotype switch in sepsis. The changes of PMN pro-inflammatory (N1 type) and anti-inflammatory (N2 type) phenotypes are studied by studying the changes of IRF4/5 subtypes, which provided a basis for identifying the infection process of PMN in clinical practice.

## Materials and Methods

### Experimental animal

The mice used in the experiments were purchased from Liaoning Changsheng Biotechnology Co., Ltd. The strains were ICR female mice, weighing 18–22g, and kept at the Animal Experimental Center of Jilin University. Animals were individually housed in a temperature-controlled (24°C) facility with 12-hour light/dark cycle (lights on 2200) with free access to food and water and allowed to acclimate for a minimum of 2 weeks after arrival. Mice were randomized according to body weight into groups. All animals were handled in accordance with the regulations established by the Animal Ethics Committee of Jilin University.

The zebrafish used in the experiment were purchased from the National Zebrafish Resource Center and raised in the Zebrafish Genetic Engineering Laboratory of Jilin Province. The feeding of zebrafish was carried out according to The Zebrafish Book and the standard operating procedure of Zebrafish Genetic Engineering Laboratory of Jilin Development and Reform Commission. Zebrafish were kept at 28.5℃ with a 14:10-h light: dark cycle in an ESEN G109 recirculating tank system using local tap water (pH 7.2–7.6, salinity 0.25±0.5‰, conductivity 500-800 µs/cm).

### Bacteria preparation

Preparation of engineered bacteria: BL21 (for zebrafish experiment) / JM109 *E·coli* strain (for mouse experiment) was inoculated in antibiotic-free LB liquid medium at a ratio of 1 ‰ −5 ‰ in an ultraclean table (220 rpm, 37 ℃ for 6 h), a small amount of bacterial fluid was taken and streaked on antibiotic-free LB solid plate, the plate was inverted, and the incubator at 37℃ overnight. a single colony of *E. coli* was picked up and cultured in 5 mL LB medium (220 rpm, 37℃ for 6 h). After 6 h of shaking culture, The *E. coli* were inoculated into Erlenmeyer flasks of LB broth in the same proportion on a sterile ultra clean table and grown by shaking for 12-16 h until the optical density (OD value) at 600 nm reached 0.7, the colony-forming units (CFUs) of the *E. coli* culture were calculated and determined to be approximately 1 × 10^8^ CFUs. For experiments involving mice, the pellet was washed twice with sterile 0.9% NaCl (saline) and then resuspended in 0.1 mL saline containing 0.1×10^8^ CFUs as a preconditioning dose and 1 mL saline containing 3×10^8^ CFUs of *E. coli* as the lethal dose, respectively. Those solutions were then ready to be used to infect the mice. For experiments involving zebrafish, the pellet was washed twice with phosphate buffered saline (PBS) and then resuspended in 0.1 mL PBS containing 0.1×10^8^ CFUs as a preconditioning dose and 1mL PBS containing 3×10^8^ CFUs of *E. coli* as the lethal dose, respectively.

Clinical pathogenic bacteria strains and culture conditions: The experimental strains were isolated and cultured from clinical infected patients in China-Japan Union Hospital of Jilin University, including *E.coli*, *Klebsiella pneumoniae* and *Staphylococcus aureus*. The frozen strains were removed from the refrigerator, inoculated in 10 mL LB liquid medium at 37℃, 225 rpm, and shaken overnight. The next day, 5 mL of overnight cultured bacterial solution was centrifuged at 13800 g for 1 min, and the bacterial precipitation was collected.

### Selection and collection of clinically isolated bacterial strains

According to the results of national bacterial resistance surveillance in 2021, the top three strains were selected, which were *E.coli*, *Klebsiella pneumoniae* and *Staphylococcus aureus*. Six strains of resistant and non-resistant *E.coli*, *Klebsiella pneumoniae* and *Staphylococcus aureus* were selected as experimental strains by collecting and analyzing the inpatients and outpatients of China-Japan Union Hospital of Jilin University, The strain numbers are as follows: non-resistant *E.coli*(ATCC25922), non-resistant *Klebsiella pneumoniae*(ATCC700603), non-resistant *Staphylococcus aureus*(ATCC25923), resistant *E.coli*(21082279), resistant *Klebsiella pneumoniae* (21086413), resistant *Staphylococcus aureus* (21082264). Blood samples were collected from patients with bacteremia. After isolation and culturing, the strains were frozen in a refrigerator at −80℃.

### Routes of delivery: exposure by immersion or microinjection

To eliminate differences arising from varying parentage, all zebrafish larvae were divided randomly into experimental and control groups. For exposure by immersion, the larvae were cultured to 3 days post fertilization (3 dpf), larvae were treated with varying concentrations of LPS by static immersion in a total volume of 100 µL per well in 96-well plates. Control larvae were exposed to regular egg water. For exposure by microinjection, prior to injection and live imaging, 3 dpf larvae were anesthetized in 0.02 % (w/v) buffered 3-aminobenzoic acid methyl ester (pH 7.0) (Tricaine; Sigma-Aldrich, A5040). The corresponding concentration of bacteria was injected into the corresponding part of the zebrafish. A negative control group was injected with PBS. For Modeling Visualization of bacterial infection in zebrafish (Result 3), The zebrafish embryos were individually infected by microinjection with 1 nL of *E·coli* either into the yolk sac space or in the total cardinal vein. For establishment of pre-adaptation model of clinical pathogenic bacteria (Figure 5E-F), Microinjection was performed with a volume of 5 nL of clinical pathogenic bacteria per larva into the yolk sac. Low-dose bacterial solution was injected for the first time with a concentration of 200 CFUs, and high-dose bacterial solution was injected for the second time with a concentration of 8000 CFUs. The interval between the two injections was 2 h. High dose group was used as control group to observe the survival rate changes of primary infection and secondary infection in 24 h. The migration and aggregation of neutrophils were observed under fluorescence microscope (Agilent BioTek Cytation 5, BioTek, Inc., Vermont, MA, USA; AE31E, motic, Inc., Xiamen, China), and the changes of neutrophils in zebrafish during primary and secondary infection were compared and analyzed by ImageJ software. All procedures involving zebrafish embryos were according to local animal welfare regulations.

### Induction of bacterial inflammation lethal model and inflammatory preconditioning model in mice

the ICR female mice weighing 18-22 g were randomly and equally divided into four groups. Blank control group (Control): mice were intraperitoneally injected with 0.1mL saline. lethal model group of sepsis (Model): mice were injected intraperitoneally with 3 × 10^8^ CFUs of *E. coli*, 1 mL/mouse. Inflammatory preconditioning group alone (InP): mice were intraperitoneally injected with prepared 0.1×10^8^ CFUs of *E·coli*, 0.1 mL/mouse. Inflammatory preconditioning protection group (InP + model, IM): mice were intraperitoneally injected with prepared 0.1×10^8^ CFUs of *E·coli*, 0.1 mL/mouse, 2 h later were injected with 3 × 10^8^ CFUs of *E. coli*, 1 mL/mouse, to establish the inflammatory preconditioning protection model of mice.

### Flow cytometry to detect of percentages of cell types

Peritoneal lavage cells were harvested in a non-enzymatic cell dissociation solution and concentrated by centrifugation from ICR female mice that had been either infected with *E. coli* or injected with saline. The dose of i.p. of mice is as described in the above. flow cytometry was used for phenotype analysis, isotype controls were routinely used in intracellular experiments. All samples were blocked with Fc-block anti-mouse CD16/32 antibody (clone: S17011E, cat. no. 156603, BioLegend). Single-cell suspensions were stained 30min at 4°C in the dark with PerCP/Cyanine5.5 anti-mouse Ly6G (clone:1A8, cat. no. 127615, BioLegend), FITC anti-mouse/human CD11b (clone:M1/70, cat. no. 101205, BioLegend) and washed three times with cold PBS. The neutrophils were selected for Ly6G^+^ and CD11b^+^ cells, according to the manufacturer’s recommendations. Neutrophils (defined as Ly6G^+^ cells with >90% purity of living cells) and non-neutrophils were obtained. After the supernatant was discarded by centrifugation, 200 μL of 4% paraformaldehyde was added and fixed overnight at room temperature (RT). the next day, after centrifugation (4°C, 1200 g, 5min), discarded the supernatant and punched with Triton X-100 (cat. no. 1139ML100, Biofroxx), add intracellular direct-labeled antibodies: PE anti-mouse IRF5 (clone: W16007B, cat. no. 158603, BioLegend) and Alexa Fluor 647 anti-IRF4(clone: IRF4.3E4, cat. no. 646407, BioLegend), incubate at 4°C in the dark for 45 min. Isotype controls (PE Rat IgG2a Isotype Ctrl, clone: RTK2758, cat. no. 400507, BioLegend; Alexa Fluor 647 sotype Ctrl, clone: RTK2071, cat. no. 400418, BioLegend) were used for each sample, and the percentage of positive staining in each sample was calculated. All flow cytometric analyses were carried out on BD FACSAria II flow cytometer (BD Biosciences) and performed with FlowJo Version 10.

### Immunofluorescence staining (IF)

Collect PLCs and centrifuge at 300 g for 10min, resuspend the cells with 100 μL PBS. All samples were blocked with Fc-block anti-mouse CD16/32 antibody (clone: S17011E, cat. no. 156603, BioLegend). After the supernatant was discarded by centrifugation, 200 μL of 4% paraformaldehyde was added and fixed overnight at RT. The next day, discard the fixative solution and wash the cells once with PBS, centrifuge (4°C, 1200 g, 5min), discard the supernatant and add 1mL Triton X-100 (cat. no. 1139ML100, Biofroxx), discard the supernatant by centrifugation, add 1 μL IRF5 antibodies (clone: 10T1, ab33478, Abcam), after samples were stained 45min in the dark at 4°C, add 1 μL PE-goat anti-mouse antibody(ab7002, Abcam), after mixing again, incubate at 4°C in the dark for 45 min. Wash the cells with PBS buffer, add 100 μL of 1 μg/mL DAPI staining solution(#4083, CST) and 2 μL CFSE staining solution (C1031, Beyotime, China) in sequence, and incubate at RT for 10 min in the dark. Then the cells were washed twice, the cell suspension was added dropwise to the center of the glass slide, 5 μL of Fluoromount-G^TM^ Anti-fade mountant (36307ES08, YEASEN) was added dropwise and covered with a cover slip, and observed with a laser confocal microscope (VHX-7000N, Keyence, Osaka, Japan).

### Fluorescent bacteria prepared by dye staining for clinical pathogens

Prepare bacteria sample with concentration in range of 10^6^ to 10^8^ cells/mL. Grow bacteria into late log phase in appropriate medium. Remove medium by centrifugation at 10,000 g for 10 min and re-suspend the pellet in Assay Buffer. Treat cells with test compounds as desired. Remove treatments by centrifugation at 10,000 g for 10 min and re-suspend the pellet in appropriate amount of Assay buffer so the concentration of bacteria in the treated sample is the same as the live. Add 1 µL of the 100X MycoLight™ 520 stock solution and 10 µL of 10X Signal Enhancer to 90 µL of the bacterial sample in Assay Buffer. Mix well and incubate in dark for 5-10 min at 37°C or 60 min at RT for optimum staining results. Monitor fluorescence of bacteria with a fluorescent microscope through FITC (Ex/Em = 488/530 nm) channel.

### Zebrafish embryo and larvae maintenance

To generate embryos, adults were placed in spawning tanks in the afternoon and then spawned following the onset of light the next day. The ratio of female and male zebrafish is 1:1 or 2:1, and the total number of zebrafish in each spawning tank is ≤4. Fertilized embryos collected after natural spawning were cultured at 28.5°C in clean Petri dishes. To collect embryos of the same period for microinjection, batches of embryos were collected every 15 min after baffling removal. Zebrafish embryos were collected within the first h post fertilization (hpf) at the same time, removed faces, whitish dead embryos and impurities and kept at 28.5 °C in E3 medium (5.0 mM NaCl, 0.17 mM KCl, 0.33 mM CaCl·2H_2_O, 0.33 mM MgCl_2_·7H_2_O) supplemented with 0.5 mg/L methylene blue (Sigma). To inhibit the formation of melanocytes, 0.003% (v/v) 1-phenyl-2-thiourea (PTU) (Sigma) was added after 10 - 12 h[42]. The larvae were cultured to 3 dpf in a periodical light incubator at 28.5°C for subsequent experiments.

### Survival experiments in infected zebrafish embryos

3 dpf zebrafish were randomly divided into groups and 10 fish per group were placed into 96 well plates. The control group was added with 100 μL of Holt buffer solution, and the other groups were added with 100 μL of the corresponding LPS solution, then the 96-well plate was incubated in a 28.5 °C constant temperature incubator in the dark for 2 h. In the establishment of a lethal model of sepsis and a protective model of inflammatory preconditioning in zebrafish larvae treated with LPS, thirty zebrafish at 3 dpf were randomly divided into three groups, and 10 fish per group were placed into 96 well plates. The control group and Model group were added with 100 μL of Holt buffer solution, InP+Model group was added 100 μL of 10 μg/mL LPS solution, then wrap them in tinfoil to avoid light and place them in a 28.5 °C constant temperature light incubator for 2 h. Slowly and carefully remove the solution in the well with a 200 μL pipette. The control group was added with 100 μL of Holt buffer solution, Model group and InP+Model group are added with 100 μL of 100 μg/mL LPS solution, the 96-well plate was further incubated for 2 h in a constant temperature incubator protected from light. After the timing is over, add fresh Holt buffer solution to the 96-well plate again. The 96-well plate was continued to be cultured and observed for 48 h. The mortality rate was determined by monitoring live and dead embryos at fixed time points. All experiments were performed in triplicates. the survival of the zebrafish in each group was recorded and the survival curve was drawn.

### Modeling visualization of bacterial infection in zebrafish

A total of 20 zebrafish at 3 dpf were randomly divided into two groups, in the bacterial local infection visualization model, which were assigned to the local infection model group (Fig. 3G-K) and the InP + local infection model group (Fig. 3L-P). The juveniles in the local infection model group were microinjected with 1 nL of 3 times *E·coli* in the yolk sac space, and the InP+local infection model group were first soaked in 100 μL of 0.1 times *E·coli* for 2 h, and then injected 1 nL of 3 times *E·coli* in the space of the yolk sac space; in the bacterial systemic infection visualization model, which were assigned to the systemic infection model group and the InP + systemic infection model group. The juveniles in the systemic infection model group were microinjected with 1 nL of 3 times *E·coli* into the total cardinal vein, and InP+systemic infection model group were first soaked with 100 μL of 0.1 times *E·coli* for 2 h and then injected 1 nL of 3 times *E·coli* into the total cardinal vein. At the end of the injection, the changes in the number of fluorescently labeled bacteria and PMNs in the larvae were recorded by taking photos at different time points with a fluorescence microscope (Agilent BioTek Cytation 5, BioTek, Inc., Vermont, MA, USA; AE31E, motic, Inc., Xiamen, China).

### Identification of clinical pathogenic bacteria and drug sensitivity test in vitro

Matrix assisted laser desorption ionization time of flight mass spectrometry (MALDI-TOF MS, Biomeriere company, France) was used to identify the bacteria isolated from patient specimens. A single colony was selected, the sample was mixed with 1μl matrix solvent, and then placed on the surface of the target plate to dry naturally, and then tested by machine to determine the name of the bacteria. General bacterial drug sensitivity test system (Vitek 2 Compact, Biomeriere company, France) and associated drug sensitivity card were used. Some tests were performed by disk diffusion method (Kirby Bauer, KB), E-test method or microbroth dilution method to obtain the minimum inhibitory concentration (MIC).

### Construction of antimicrobial susceptibility test platform in zebrafish

Eight antibiotics including meropenem, cefepime, levofloxacin, ceftazidime, amikacin, piperacillin-tazobactam, penicillin and benzoxicillin (The above reagents were purchased from Sigma-Aldrich) were selected for drug toxicity test in zebrafish using automatic bacterial identification. Concentrations of all antibiotics were based on the MIC value of the antibiotic for each strain (Supplementary Table S4). After the gradient dilution of antibiotics with PBS, the larvae were soaked in the drug solution for 24 h to observe the toxicity of the drug to the zebrafish, and to select a safe drug concentration that had no effect on the morphology and behavior of the zebrafish at 24 h for drug screening in vivo.

For the establishment of clinical pathogenic bacteria models, microinjection was performed with a volume of 5 nL of clinical pathogenic bacteria per larva into the yolk sac space. 3 dpf zebrafish were injected with six fluorescent strains respectively and soaked for 2 h with antibiotics with different MIC multiple concentrations in the above-mentioned different commonly used clinical antibiotic treatment schemes respectively. After soaking for 2 h, they were kept in new regular egg water, and the changes of bacterial fluorescence intensity in zebrafish were observed under fluorescence microscope at 0, 2 and 4 h, and were analyzed by ImageJ software after taking photos. At the same time, the survival rate of zebrafish in 24 h was observed after treatment with antibiotics with different concentrations of MIC. Transgenic zebrafish with red neutrophil fluorescence labeled and green neutrophil fluorescence labeled respectively named Tg (LysC: DsRed2) and Tg (mpx: EGFP) were used for infection experiments to observe the interaction between neutrophil and host and the aggregation and migration of neutrophil during infection. ImageJ software was used to analyze the changes in the number distribution of neutrophil in the anterior and tail of zebrafish for 0∼6 h.

### RNA extraction and real-time quantitative PCR (qPCR) analysis

A total of 30 zebrafish larvae at 3 dpf were placed in 1.5 mL EP tubes, and 1 mL trizol was added after removing the Holt buffer solution. Total RNAs were harvested by using the trizol reagent per the manufacturer’s instructions (Invitrogen, Carlsbad, CA, USA). cDNA synthesis was performed using the Super RT Kit (BioTeke, Beijing, China) according to the manufacturer’s protocol. qPCR experiments were conducted using a 2× plus SYBR real-time PCR mixture (BioTeke, Beijing, China) according to the manufacturer’s instructions. The qRT-PCR experiment was done with an ABI 7500 Real Time PCR System (Applied Biosystems, Carlsbad, CA). Each cDNA sample of each target gene (TGF-β, IL-1β, IL-10, IL-8, IRF5, IRF4, TNF-α) was detected in triplicate, and the results were normalized to the expression level of *β*-actin (ACTB) and expressed relative to the control group. the relative expression levels were calculated with the formula 2^−ΔΔCt^. All primer designs were obtained by Primer 5 software (Premier Corporation, Canada) and purchased from Sangon Biotech (Sangon, Shanghai). The following primers were used to detect the expression level of *TGF-β1a,* forward: 5’-TGAGGCTATTCGGGGTCAGA-3’and reverse:5’- GGAGACAAAGCGAGTTCCCA-3’; *IL-1β,* forward: 5’- GGACTTCGCAGCACAAAATGAA-3’ and reverse: 5’- TTCACTTCACGCTCTTGGATGA-3’; *IL-10,* forward: 5’- AGCACTCCACAACCCCAATC-3’ and reverse: 5’- AGCAAATCAAGCTCCCCCATA-3’; *IL-8,* forward: 5’- GGAGATCTGTCTGGACCCCT-3’ and reverse: 5’- TGATCCGGGCATTCATGGTT-3’; *IRF-5,* forward: 5’- TAGTGGACAGCCCAATGCAG-3’ and reverse: 5’- TCCAAAATCAGCCCTCGGTC-3’; *IRF-4,* forward: 5’- TCGCCATCAGAAAGTCACCC-3’ and reverse: 5’- GCGACCGTGATGAATGAAGC-3’; *TNF-α,* forward: 5’- TGCAAAACTCGGCCAATTCC-3’ and reverse: 5’- CCCGAAGAATGTTTTGGCGT-3’, and the primer sequences for the control gene *β-actin* were 5’-ACCACGGCCGAAAGAGAAAT-3’ (Forward) and 5’- ATGTCCACGTCGCACTTCAT-3’ (Reverse).

### Western blotting

Western blotting was done using standard protocol. Briefly, total protein was prepared using RIPA Lysis Buffer (BostonBio Products, Ashland, MA, USA) containing PMSF (Sigma) and quantified with a BCA Protein Assay Kit (Beyotime, China). According to the protein concentration, a buffer solution was added, and the mixture was boiled for 10 min. Same amounts of proteins were separated on a 10% SDS-polyacrylamide gel, and transferred onto a nitrocellulose membrane. The nitrocellulose membrane was incubated with 5% non-fat milk in Tris-buffered saline (150 mM NaCl, 20 mM Tris-HCl, pH7.4) with primary antibody overnight at 4°C. After washing, the membrane was further incubated with secondary antibody for 45 min and proteins were detected with an ECL detection system. Image J software was used to measure the band intensity.

### Co-Immunoprecipitation experiment (Co-IP)

Immunoprecipitation studies were done as described previously [43]. Peritoneal lavage cells were isolated, homogenized, lysed, and centrifuged, and the supernatant was collected afterward. After a 2 h incubation at 4°C, the complex containing supernatant and antibody against IRF4, IRF5 or MyD88 was mixed with r-protein-G agarose. The beads were washed four times with solubilization buffer to remove the bound proteins and heated at 95°C for 5 min in SDS sample buffer. Samples were separated by SDS-polyacrylamide gels and then transferred onto nitrocellulose membranes, blocked with 5% milk, incubated with primary antibodies and secondary antibodies, then measured with an ECL detection system.

### Morphological observations of zebrafish

Use a plastic straw to suck out the larvae and make them lie on their sides in a clean dish, try to make the eyes of the larvae overlap, and the body is in a natural horizontal state. By translating the dish in different directions, the juveniles were located in the center of the field of view, and then the microscope was adjusted to make the field of view clear. At this time, the morphological changes of individual juveniles in each group were recorded by photographing software. The main morphological changes were as follows: Tail damage and shrinkage gradually increased; the curvature of the spine increases gradually; the body necrosis gradually increased, and the body color of the larva changed from transparent to black; in severe cases, the yolk sac of the larva is damaged and the content flows out.

### Imaging and fluorescence quantification

A fluorescence stereo microscope (SZX10, Olympus, Inc., Tokyo, Japan) was used to take images of zebrafish larvae. During imaging, larvae were kept under anesthesia with 0.02% Tricaine (Sigma). To quantify fluorescence of bacterial burden in individual larvae, the fluorescent images of larvae with custom-made, dedicated pixel quantification software were performed.

### Statistical analysis

Statistical analysis was performed using GraphPad Prism 8 (GraphPad Software, La Jolla, CA, USA). Survival experiments were evaluated using the Kaplan–Meier method. Comparisons between curves were made using the log rank test. Differences in bacterial burden were statistically tested by one-way ANOVA followed by Tukey’s comparison test (multiple group comparisons). For qRT-PCR, statistical significance among multiple groups was estimated ANOVA on ln(n)-transformed relative induction folds. Significance (*P*-value) is indicated with ns, non-significant; * *P* < 0.05; ** *P* < 0.01; *** *P* < 0.001. All experiments were performed three times and Error bars are the mean ± standard error mean (SEM).

## Ethics declarations

All mouse studies were approved by the Institutional Animal Care and Use Committee (IACUC) of Jilin University, approval number is SYXK(Ji) 2013-0005. The blood samples of patients were used in this research provided written informed consent. The study protocol was approved by the Clinical Research Ethics Committee of China-Japan Union Hospital of Jilin University.

